# The pentatricopeptide repeat protein MTSF3 is required for *nad2* mRNA stability and embryogenesis in Arabidopsis

**DOI:** 10.1101/2022.05.20.492872

**Authors:** Chuande Wang, Lisa Blondel, Martine Quadrado, Céline Dargel-Graffin, Hakim Mireau

**Affiliations:** Université Paris-Saclay, INRAE, AgroParisTech, Institut Jean-Pierre Bourgin (IJPB), 78000, Versailles, France

## Abstract

Gene expression in plant mitochondria is predominantly governed at the post-transcriptional level and relies mostly on nuclear-encoded proteins. However, the involved protein factors and the underlying molecular mechanisms are still not well understood. In this study, we report the function of the mitochondrial stability factor 3 (MTSF3) protein and we show that it is essential for accumulation of the mitochondrial *nad2* transcript in Arabidopsis and not for the splicing of *nad2* intron 2, as recently proposed (Marchetti et al., 2020). The MTSF3 gene encodes a pentatricopeptide repeat protein that localizes in the mitochondrion. An *MTSF3* null mutation induces embryonic lethality but viable *mtsf3* mutant plants could be generated by partial complementation with the developmentally-regulated *ABSCISIC ACID INSENSITIVE3* promoter. Genetic analyses reveal that *mtsf3* rescued plants display growth retardation due to the specific destabilization of a *nad2* precursor transcript bearing exons 3 to 5. Biochemical data demonstrate that MTSF3 protein specifically binds to the 3’-terminus of *nad2*. The destabilization of *nad2* mRNA induces a significant decrease in complex I assembly and activity, and an overexpression of the alternative respiratory pathway. Our results support that the MTSF3 protein protects *nad2* transcript from degradation by mitochondrial exoribonucleases by binding to its 3’ extremity.

## INTRODUCTION

Mitochondria and chloroplasts are two cellular organelles that descend from free-living bacterial ancestors and were acquired through endosymbiosis (Zimorski et al., 2014). During subsequent host-endosymbiont co-evolution, both organelles lost most genes inherited from their bacterial ancestors and have retained limited sets of genes that are essential for their activity (Timmis et al., 2004). Unlike bacteria, in which gene expression is primarily regulated at the transcriptional level, modern organelles possess a surprisingly complex mRNA metabolism involving numerous post-transcriptional RNA modifications playing predominant roles in organellar gene regulation. Post-transcriptional processing of organellar RNAs include C-to-U RNA editing, *cis*- and *trans*-splicing of group I and group II introns, 5’- and 3’- end RNA maturation as well as mRNA translation. These processes rely almost exclusively on nuclear-encoded RNA binding proteins that were acquired during nucleus-organelle co-evolution (Stern et al., 2010; Germain et al., 2013; Hammani and Giege, 2014). The underlying mechanisms remain poorly understood at the molecular level, notably in plant organelles, and the vast majority of involved protein cofactors have not been identified yet. However, over the last decade or so, several classes of RNA-binding proteins were found to play essential roles in organellar gene expression. The most represented type concerns the pentatricopeptide repeat (PPR) proteins comprising 2 to 30 repeats of highly degenerate 31–36 amino acid motifs (Small and Peeters, 2000; Barkan and Small, 2014). PPR proteins constitute a large family of RNA binding proteins that dramatically expanded in terrestrial plants, with more than 450 members in Arabidopsis (Lurin et al., 2004; O’Toole et al., 2008). PPR proteins can be classified into two major groups based on the nature of their motifs with P-type proteins containing only canonical 35 amino-acid motifs and PLS-type proteins possessing triplets of P, L (Long, 35 or 36 amino acids) and S (Short, 31 or 32 amino acids) motifs along with carboxy-terminal domains named E1, E2 or DYW (Lurin et al., 2004; Cheng et al., 2016). The P-class PPR proteins participate in various aspects of organellar RNA expression ranging from gene transcription to mRNA translation, while PLS-PPR proteins are almost exclusively implicated in C-to-U RNA editing (reviewed in (Barkan and Small, 2014; Gorchs Rovira and Smith, 2019; Small et al., 2020)). Structural studies have revealed that PPR proteins exhibit a right-handed superhelical α-solenoid structure that consists of successive tandem repeats, each of which forming a pair of antiparallel α-helices (Yin et al., 2013; Shen et al., 2016). The recognition between the PPR proteins and their RNA ligands involves a one-motif to one-base rule, in which combinations of two residues at positions 5 and 35 in each repeat are responsible for specific RNA base recognition (Barkan et al., 2012; Takenaka et al., 2013; Yagi et al., 2013; Yan et al., 2019).

Before being translated, plant mitochondrial mRNAs most often require 5’ and 3’ shortening to become mature. Several PPR proteins, called RNA processing factors (RPFs), have been found to participate in 5’-end processing of Arabidopsis mitochondrial transcripts (reviewed in (Binder et al., 2016)). Most RPFs were classifi ed as restorer of fertility-like (RF-like or RFL) proteins because of their strong similarity with restorers of fertility (Rf) proteins identified in various crop species (Jonietz et al., 2010; Hölzle et al., 2011; Jonietz et al., 2011; Arnal et al., 2014; Stoll et al., 2017; Schleicher and Binder, 2021). However, 5’-end processing seems not always crucial for the functioning of plant mitochondrial mRNAs, as most *rpf* mutants accumulate incorrect but equally stable 5’ extended mRNAs that are correctly translated. It is also currently unclear how these RPF-PPR proteins act to define mitochondrial mRNA 5’ ends but the recruitment of a yet unknown endoribonuclease has been suggested (Stoll and Binder, 2016). In contrast, only two PPR proteins involved in the 3’-end processing of the mitochondrial transcripts were reported so far (Haili et al., 2013; Wang et al., 2017). Mitochondrial stability factor 1 (MTSF1) and MTSF2 from Arabidopsis have been shown to be both required for the 3’-end maturation and stability of mature *nad4* mRNA and *nad1* exon 2-3 precursor RNA, respectively, by associating with the 3′ end of corresponding mitochondrial transcripts. It has been proposed that the PPR proteins stabilizes mRNAs by blocking the progression of 3’-to-5’ exoribonucleases along transcripts, which leads their binding site to correspond to the 3’ end of the RNAs they stabilize. In the case of MTSF1, the stabilized 3’ extremity corresponds to that of the mature *nad4* mRNA whereas, for MTSF2, it corresponds the 3′ end of the first half of a *trans*-intron found at the end of a *nad1* precursor transcript and thereby defines a 3’ extremity within a half intron that is compatible with a *trans*-splicing reaction needed to produce a mature mitochondrial mRNA. The range of transcripts stabilized by association with protective protein on their 3′ end is thus not limited to the 3’ extremity of mature mRNAs and can be more diverse than initially expected. In this study, we clarify the function of a mitochondria-targeted PPR protein in Arabidopsis that we named mitochondria stability factor 3 (MTSF3) protein. Effectively, this PPR protein was inconclusively characterized in a previous study (Marchetti et al., 2020) and no firm function could be assigned to it. Our biochemical and genetic analyses demonstrate that MTSF3 indeed associates with high affinity to an RNA region corresponding to the 3’ end of mature *nad2* mRNA and that of a *nad2* precursor transcript. We further provide evidence that MTSF3 protein is required for the 3’ end processing and stability of these two *nad2* transcripts. We also reveal that *mtsf3* null mutants are embryo-lethal, unlike most characterized Arabidopsis respiratory complex I mutants.

## RESULTS

### MTSF3 encodes a PPR-P protein required for embryo-development

The *mtsf3* mutant (SAIL_359_F11) was originally identified in a series of Arabidopsis T-DNA mutants having insertions in nuclear genes encoding mitochondria-targeted PPR proteins. The corresponding PPR gene corresponded to AT2G02150 gene, which was previously named EMBRYO DEFECTIVE 2794 (Meinke et al., 2008). This protein is predicted to comprise 16 canonical P-type PPR repeats according to the PlantPPR database (http://ppr.plantenergy.uwa.edu.au/). We requalified EMB2794 as MTSF3 (Mitochondrial Stability Factor 3) after molecular characterization of its function in mitochondrial RNA processing (see below). The T-DNA insertion (SAIL_359_F11) in *MTSF3* locates in the sequence encoding the sixth PPR repeat, precisely 1,082 nucleotides downstream of the start codon (Figure 1). No homozygous mutants could be recovered in the progeny of selfed *mtsf3* heterozygous plants, suggesting that *mtsf3* could be essential for embryo development. Segregation analysis further revealed that about a quarter of seeds recovered from self-pollinated heterozygous plants could not germinate, suggesting a germination deficiency associated with a single recessive mutation. Accordingly, immature siliques from heterozygous *MTSF3/mtsf3* plants contained about one-quarter of white translucent seeds (Figure 1B). To determine whether embryonic development in these seeds was perturbed, embryos from white and green seeds were examined using differential interference contrast microscopy. The results indicated that while embryos from green seeds contained fully-developed embryos, embryos from translucent seeds did not mature beyond the heart-torpedo transition stage (Figure 1C).

**Figure 1.**
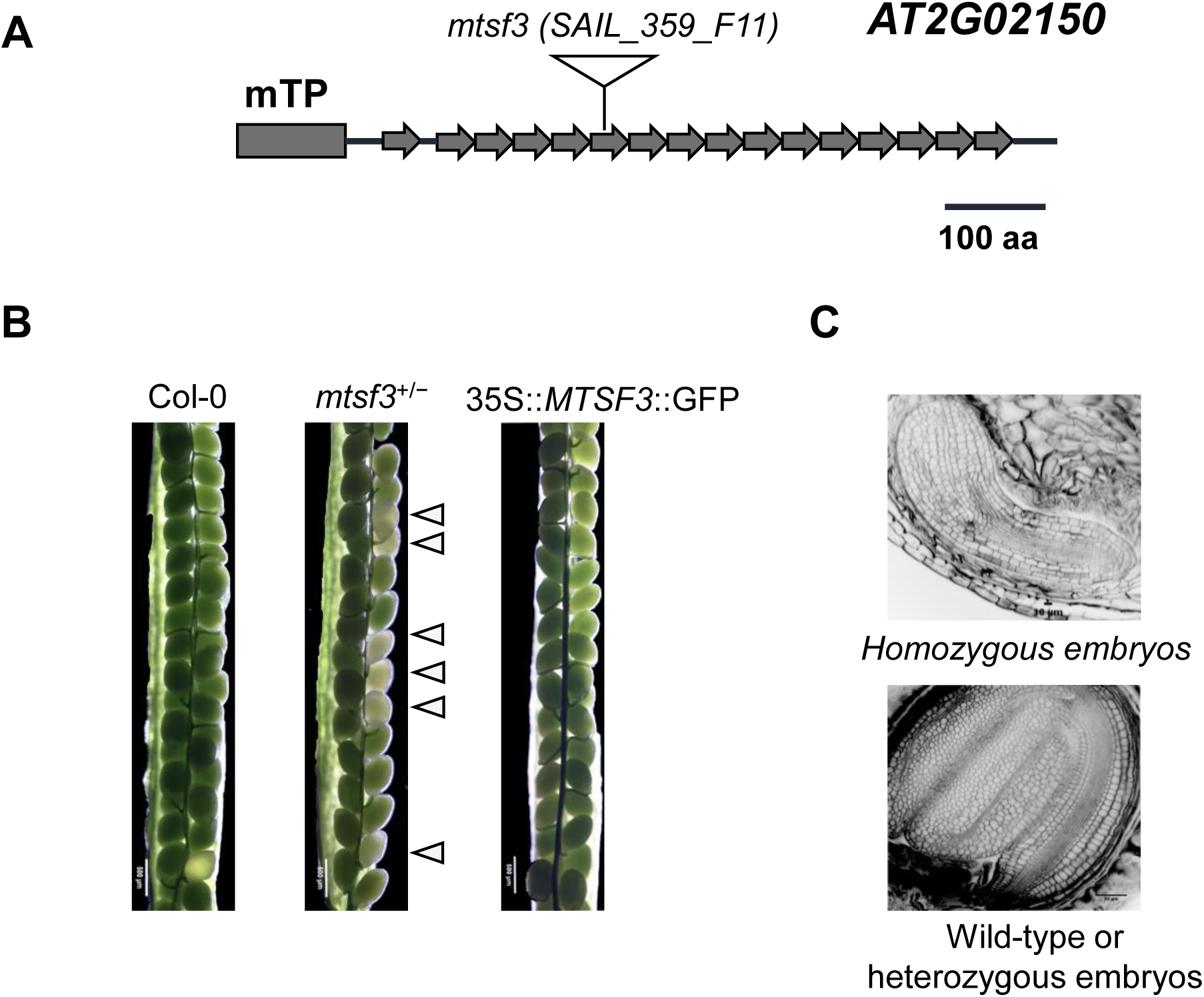
A null allele of the Arabidopsis *MTSF3* gene confers embryo lethality. (A) Schematic diagram of the *MTSF3* gene structure with the position of the SAIL_359_F11 T-DNA insertion. The length of the putative mitochondrial targeting sequence (mTP) was predicted using TargetP and is shown as a black box. (B) Open siliques showing the seeds produced from wild-type (Col-0), heterozygous and functionally-complemented *mtsf3* plants. Arrows point towards white seeds that are observed in siliques from *mtsf3* heterozygous plants. Bars=50 μm. (C) Confocal images of embryos dissected from immature seeds (6 weeks after sowing) showing the phenotype of *mtsf3* mutant embryos compared to an embryo contained in a green seed.

To confirm that the observed embryo-defective phenotype was caused by the T-DNA insertion in the *MTSF3* locus, *mtsf3* heterozygous mutants were transformed with a DNA construct comprising the AT2G02150 gene fused to the GFP under the control of the 35S promoter (35S::AT2G02150::GFP). The resulting transgenic *mtsf3* homozygous mutants displayed a restored wild-type phenotype (Figures 1 and 2), supporting that the inactivation of the AT2G02150 gene was responsible for the embryonic lethal phenotype of *mtsf3* mutant.

**Figure 2.**
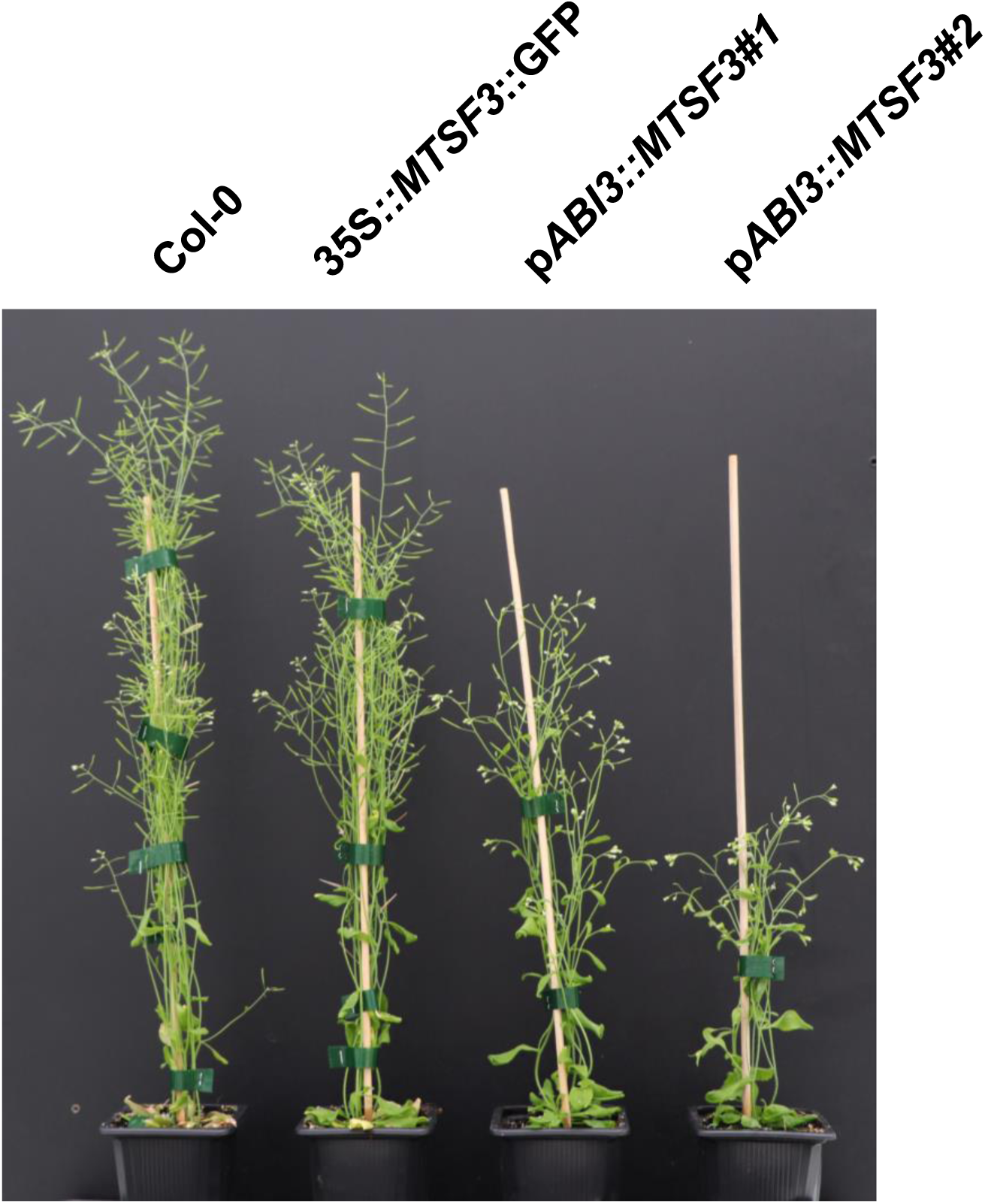
Partially-complemented p*ABI3*::*MTSF3* plants display a globally retarded growth phenotype. Photograph of five-week-old plants showing the reduced size of two partially-complemented (p*ABI3*::*MTSF3*#1 and #2) *mtsf3* mutant plants as compared with the wild type (Col-0) and a fully-complemented (35S::*MTSF3*::*GFP*) *mtsf3* homozygous mutant.

### Partially-complemented *mtsf3* mutants display a globally retarded growth phenotype

To obtain sufficient mutant material for the characterization of the MTSF3 protein, we produced partially-complemented *mtsf3* plants by expressing the *MTSF3* gene under the control of the *ABSCISIC ACID-INSENSITIVE3* (*ABI3*) promoter (p*ABI3*::*MTSF3*) (Figure 2). The *ABI3* promoter is a seed specific promoter that was used to direct the expression of *MTSF3* in *mtsf3* homozygous mutants during embryogenesis and then turn off *MTSF3* production in latter developmental stages, as previously employed for the analysis of several other embryonic lethal Arabidopsis genes (Despres et al., 2001). Two independent partially-complemented lines (p*ABI3*::*MTSF3*#1 and p*ABI3*::*MTSF3*#2) were obtained this way and used for further experimental analyses (Figure 2). Under conventional greenhouse conditions, both lines displayed a marked slow growth phenotype and harbored small twisted leaves. Additionally, the p*ABI3*::*MTSF3*#1 line exhibited a milder retarded phenotype compared with the p*ABI3*::*MTSF3*#2 line (Figure 2).

### MTSF3 is a mitochondria-localized protein

According to the Arabidopsis subcellular database SUBA (Hooper et al., 2017) and the TargetP 2.0 prediction program (Armenteros et al., 2019), the MTSF3 protein was strongly suspected to be addressed to Arabidopsis mitochondria. To verify the cellular distribution of MTSF3, a GFP C-terminal translational fusion comprising the complete sequence of *MTSF3* was generated and transformed into the *Arabidopsis* cell suspension culture PSB-D (Van Leene et al., 2011). The GFP fluorescent signal was detected to co-localize with the red fluorescence of the MitoTracker™ control, confirming the mitochondrial localization of the MTSF3 protein (Supplemental Figure S1).

### The accumulation of complex I is impaired in p*ABI3*::*MTSF3* plants

The mitochondrial localization of the MTSF3 protein suggested that the developmental defects in p*ABI3*::*MTSF3* plants could result from altered respiration. To clarify the origin of this potential respiratory deficiency, we examined the levels of the different respiratory complexes in comparison with the wild type, by blue native polyacrylamide gel electrophoresis (BN-PAGE). Mitochondria-enriched pellets were prepared from both wild-type and partially-complemented *mtsf3* plants and solubilized in 1% n-dodecyl β-D-maltoside. Following gel migration, the abundance of respiratory complexes was either visualized by in-gel activity staining or by western blot analysis with appropriate antibodies. Compared to wild-type, most respiratory complexes accumulated to normal or slightly higher levels in both p*ABI3*::*MTSF3* lines (Supplemental Figure S2). In-gel activity staining however revealed that the abundance of complex I was severely reduced in p*ABI3*::*MTSF3* plants, which was further confirmed by immunodetection with an antibody against a complex I subunit, the carbonic anhydrase 2 (Figure 3A). Higher levels of complex I could be detected in p*ABI3*::*MTSF3#1* plants compared to p*ABI3*::*MTSF3*#2 (Figure 3), which correlated with the milder growth alteration of p*ABI3*::*MTSF3#1* mutants compared to p*ABI3*::*MTSF3*#2 plants (Figure 2).

**Figure 3.**
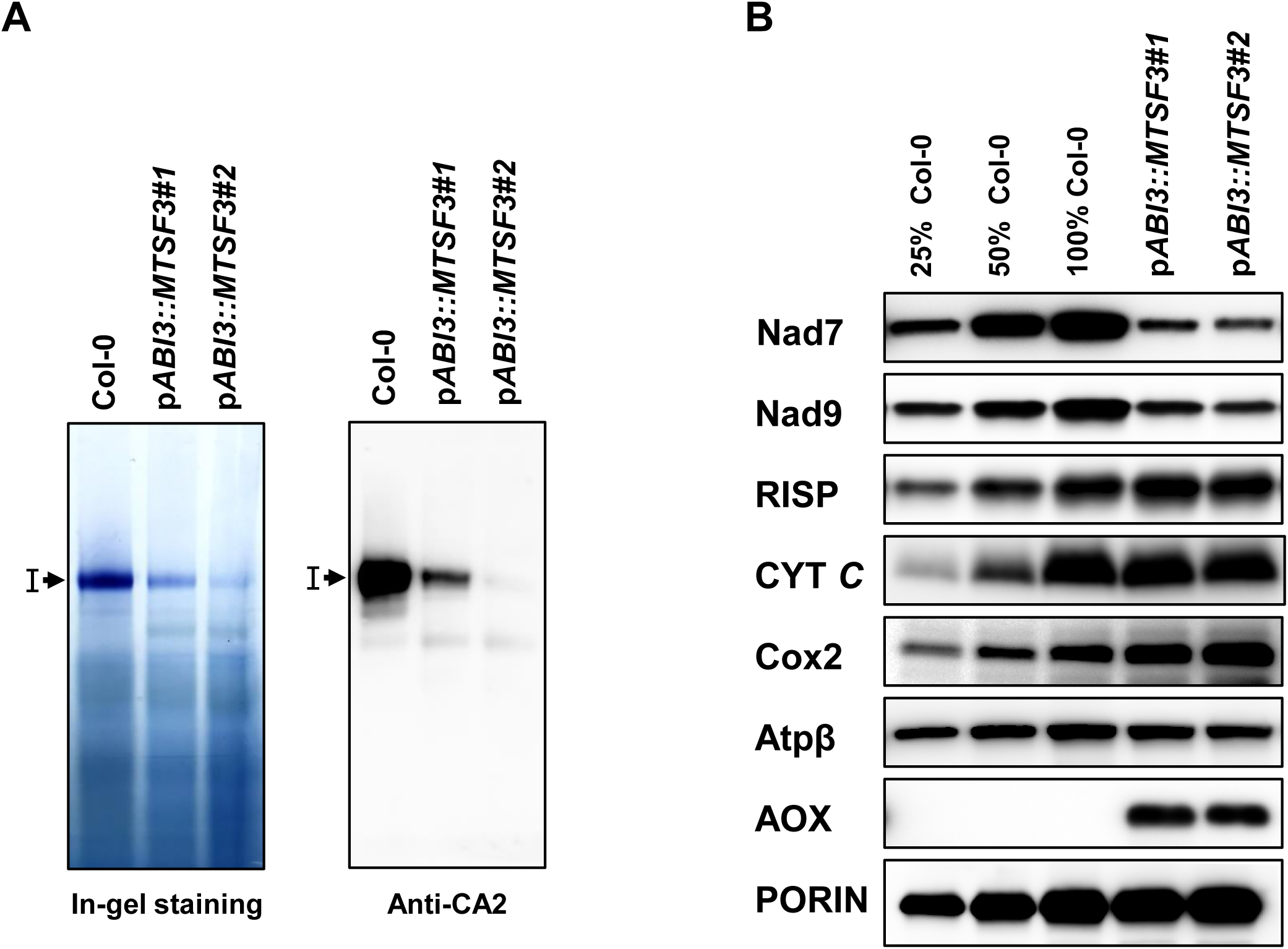
Respiratory complex I is decreased in partially-complemented *mtsf3* plants. (A) BN-PAGE analysis of mitochondrial complex I in p*ABI3*::*MTSF3* plants. In-gel activity staining revealing the NADH dehydrogenase activity of complex I is presented in the left panel. Detection of complex I was also performed on BN-PAGE blots with antibodies to the mitochondrial CA2 (carbonic anhydrase 2) subunit as shown on the right panel. Holocomplex I is indicated by an arrowhead. (B) Analysis of mitochondrial proteins steady-state levels. Crude membrane extracts from the indicated genotypes were separated by SDS–PAGE and probed with antibodies to subunits of complex I (Nad7 and Nad9), complex III (RISP), complex IV (Cox2), the ATP synthase (ATPβ), cytochrome *c* (CYT *C*) and the alterative oxidase (AOX). Porin was used as protein loading control. Dilution series of proteins extracted from the wild type (Col-0) were used for signal comparison.

We next analyzed the steady-state levels of various subunits of mitochondrial respiratory complexes by immunoblot analysis. The results showed that the abundance of RISP (a subunit of complex III), CYT*C* (cytochrome *c*, a mobile protein between complex III and IV), Cox2 (a subunit of complex IV) and ATP-β (a subunit of complex V) proteins were higher or close to normal levels in both p*ABI3*::*MTSF3* plants as compared with the wild-type (Figure 3B). These observations are consistent with corresponding respiratory complex abundances observed in BN-PAGE gel analysis (Supplemental Figure S2). In agreement with the dramatic reduction in complex I, the accumulation of Nad7 and Nad9 proteins (two subunits of complex I) was greatly reduced in the two p*ABI3* partially-complemented plants (Figure 3B).

To further characterize the respiratory activity in p*ABI3*::*MTSF3* plants, we assessed the induction of the alternative respiratory pathways by measuring the expression levels of alternative NADH dehydrogenases (*NDA, NDB* and *NDC*) and alternative oxidase (*AOX*) genes. Quantitative RT-PCR indicated that the *NDA2* and *NDB4* mRNAs over-accumulated respectively 4 and 16 times and two of the *AOX1* transcripts accumulated respectively four and six times more in p*ABI3*::*MTSF3* plants compared to the wild type (Supplemental Figure S3). The overexpression of *AOX* was further estimated by probing total mitochondrial protein extracts with an antibody to AOX1 (Figure 3B), which confirmed the overaccumulation of the alternative oxidases in the mutants. The activation of the alternative respiratory pathways was consistent with what had been previously observed in many complex I mutants (de Longevialle et al., 2007; Keren et al., 2012; Haili et al., 2013; Fromm et al., 2016; Haili et al., 2016; Wang et al., 2017; Wang et al., 2018; Wang et al., 2020). Altogether, our results suggested that the biogenesis and the activity of mitochondrial complex I was severely compromised in the p*ABI3*::*MTSF3* plants and that this resulted in a strong activation of the alternative respiratory pathway.

### MTSF3 is required for the accumulation of the *nad2* exon 3 – 5 precursor transcript

Previous studies have implicated PPR proteins in a wide range of RNA processing events in plant organelles (Barkan and Small, 2014; Hammani and Giege, 2014). The complex I deficiency in p*ABI3*::*MTSF3* plants prompted us to consider that the MTSF3 protein could play a key role in the processing of one or several mitochondria-encoded mRNAs encoding a complex I subunit. To test this hypothesis, the steady-state levels of mature and precursor mitochondrial transcripts were measured by quantitative RT-PCR in p*ABI3*::*MTSF3* and wild-type plants (Figure 4). Although the majority of mature mitochondrial mRNA accumulated at the same or only slightly higher levels in the mutants, we could observe that the level of several *nad2* transcripts were severely reduced in the two p*ABI3*::*MTSF3* lines compared to wild-type plants. The Arabidopsis *nad2* gene comprises five exons that are distributed in two distant genomic locations. Maturation of *nad2* mRNAs requires thus the fusion of two distinct pre-mRNAs named *nad2* exon 1–2 and *nad2* exon 3–5 that are reunified by a *trans*-splicing reaction involving the two halves of *nad2* intron 2, named 2a and 2b (Figure 5A). To better understand the defect in *nad2* expression in p*ABI3*::*MTSF3* plants, the steady levels of the two precursor transcripts were monitored by quantitative RT-PCR at different positions. This revealed a slight over-accumulation of pre-mRNAs containing non-spliced intron 2a, whereas precursors bearing other introns, especially introns 2b, 3 and 4, slightly down-accumulate in p*ABI3*::*MTSF3* plants (Figure 5B). Therefore, the reduction of mature transcripts spliced for introns 3 and 4 (PCR nad2ex34 and nad2ex45) are not accompanied by an increased accumulation of corresponding unspliced precursors (int3-ex4 and int4-ex4) as it always occurs in true mitochondrial intron splicing mutants (Kuhn et al., 2011; Francs-Small et al., 2012; Hsieh et al., 2015; Weissenberger et al., 2017; Wang et al., 2018; Wang et al., 2020). Such lack of inverted correlation between the abundance of precursor and mature *nad2* transcripts in *mtsf3* mutants suggested that the loss of mature *nad2* could rather result from an instability of *nad2* exon 3-5 pre-mRNAs rather than a splicing deficiency of a *nad2* intron *per se*. As the *nad2* exon 3-5 pre-mRNA is engaged in a *trans*-splicing reaction involving intron 2, its instability inevitably affects the splicing of *nad2* intron 2 manifested by an increase of precursors containing intron 2a (PCR int2a-ex2 of Figure 5B) and a decrease in mature *nad2* mRNAs spliced for intron 2 (PCR nad2ex23 of Figure 4B). RNA gel blots were then performed to validate our qRT-PCR results (Figure 5D). Coherently, only trace amounts of mature *nad2* mRNA and *nad2* exon 3-5 precursor form could be detected in the p*ABI3*::*MTSF3* plants. The slight over-accumulation of *nad2* exon 1-2 precursor transcript detected by quantitative RT-PCR was also visible in RNA gel blots.

**Figure 4.**
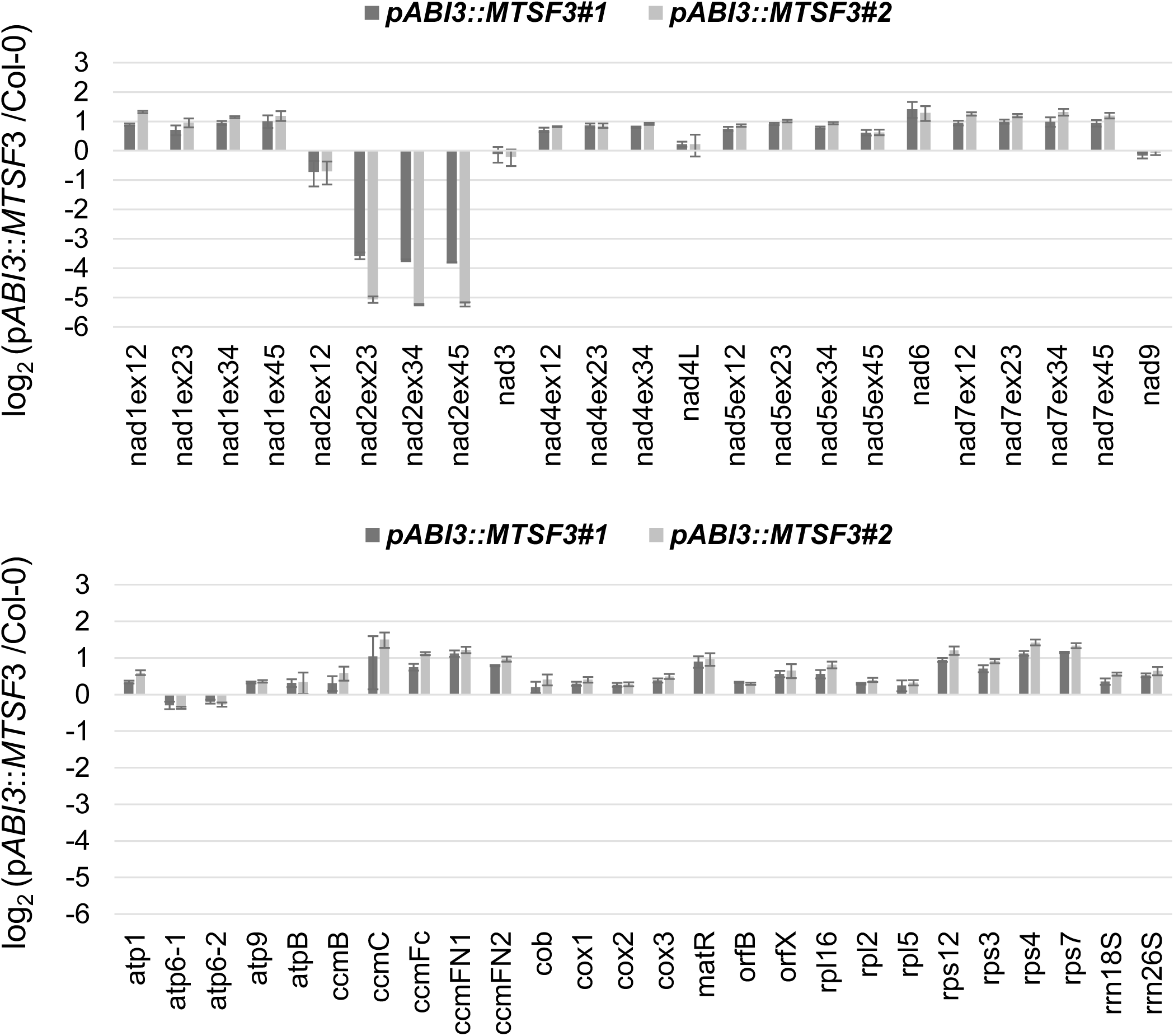
Mature *nad2* mRNA strongly under-accumulate in partially-complemented *mtsf3* mutants. The steady state levels of mature mitochondrial mRNAs measured by quantitative RT-PCR in Col-0 and partially-complemented *mtsf3* plants. The histograms show log_2_ ratios of p*ABI3*::*MTSF3* to wild-type. A single PCR was considered for mRNAs carrying no introns, whereas the accumulation of individual exons was analyzed for intron-containing transcripts. Three biological replicates and three technical replicates were used per genotype; standard errors are indicated. The data were normalized to the nuclear 18S rRNA gene.

**Figure 5.**
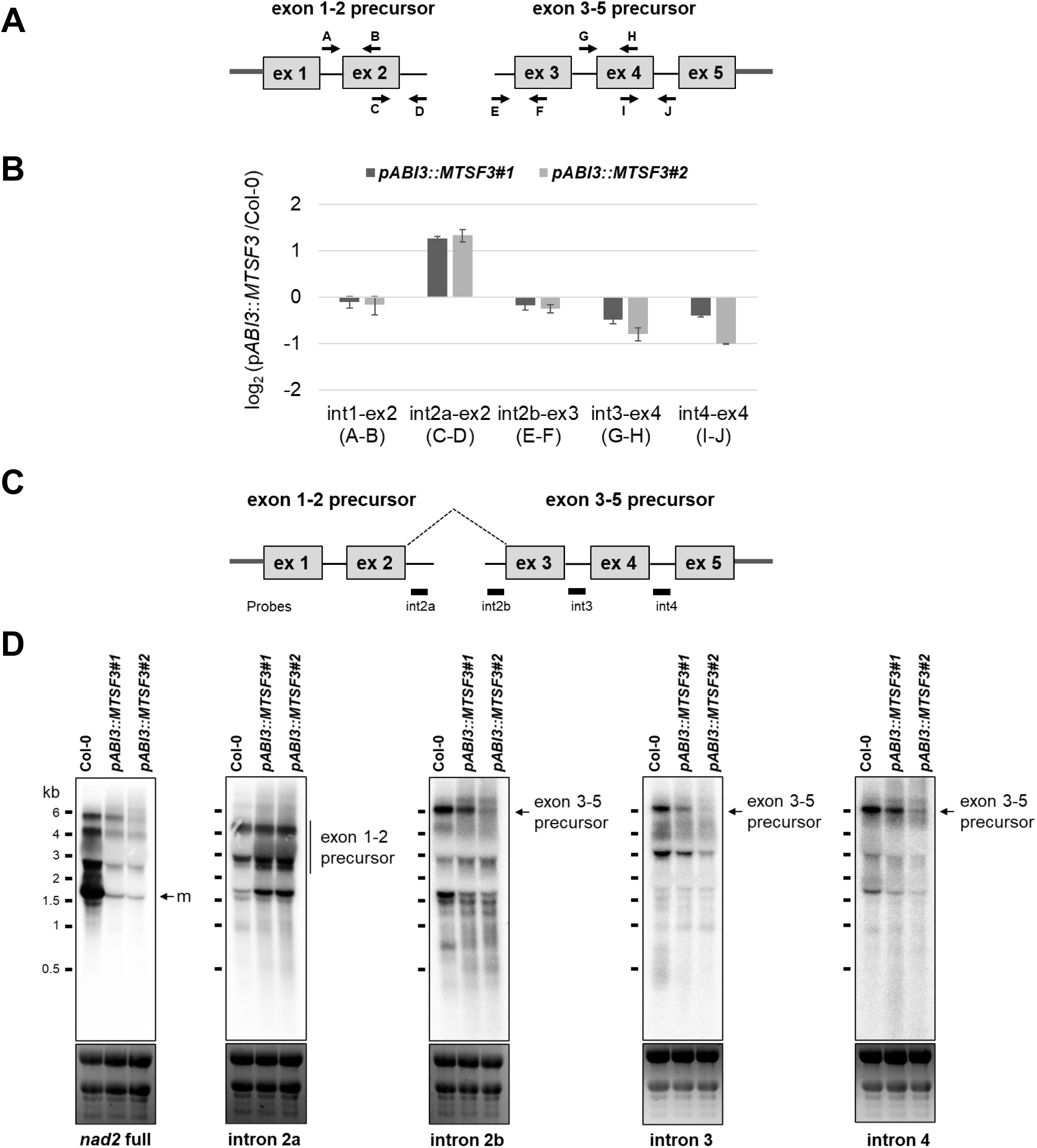
The stability of the *nad2* exon 3–5 precursor is compromised in partially-complemented *mtsf3* mutants. (A) Schematic representation of *nad2* exon 1-2 and *nad2* exon 3–5 precursor transcripts and positions of the primers used for quantitative RT-PCR analysis. (B) Quantitative RT-PCR measuring the steady state levels of *nad2* exon 1-2 and *nad2* exon 3–5 precursor RNAs in Col-0 and partially-complemented *mtsf3* plants. The histograms show log_2_ ratios of p*ABI3*::*MTSF3* plants to wild-type. The PCR reactions were performed with the indicated primer pairs. Three biological replicates and three technical replicates were used per genotype; standard errors are indicated. (C) Schematic representation of the two *nad2* precursor transcripts, which are fused by one *trans*-splicing event to produce the mature (m) *nad2* mRNA. Gray boxes indicate exons. Introns and 5′ and 3′ UTRs are shown as thick and thin lines, respectively. The probes used to interpret RNA gel blot results are also indicated. (D) RNA gel blots showing the accumulation profiles of *nad2* transcripts in the two partially-complemented *mtsf3* mutants compared to WT (Col-0). Used probes are indicated below hybridization results. Ethidium bromide staining of ribosomal RNAs is shown below the blots and serves as a loading control. m: mature mRNAs.

Overall, these results strongly suggested that the loss of mature *nad2* mRNA results from a specific destabilization of *nad2* exon 3–5 precursor transcript in p*ABI3*::*MTSF3* plants, and thus that the MTSF3 protein plays likely a role in the stabilization of this precursor mRNA.

### MSTF3 specifically associates with the 3’ region of *nad2 in vivo* and *in vitro*

To further dissect the possible role of MTSF3 in the stabilization of *nad2* mRNA, the putative RNA binding site of MTSF3 was predicted according to the previously established PPR recognition code (Barkan et al., 2012). The predicted RNA recognition sequence was then used to scan the Arabidopsis mitochondrial genome (Figure 6A). Of the top ten predicted binding sites, two concerned the *nad2* gene, with one possible binding site in the first half of intron 2 and the other in the 3’ UTR region (Figure 6B). To test whether the MTSF3 associates with these RNA targets or not, RNA immunoprecipitation assays followed by reverse transcription and quantitative PCR (RIP-RT-qPCR) assays were used. To this end, an Arabidopsis transgenic cell line expressing the MTSF3 protein in fusion with the GFP tag was first produced. The MTSF3-GFP fusion was then immunoprecipitated from transgenic cell extracts and coenriched RNAs purified from the coimmunoprecipitates and used for RT-qPCR analysis (Figure 6C). A highly specific co-enrichment of MTSF3 with the 3’ UTR of *nad2* was revealed, supporting an association of MTSF3 with this *nad2* region. In contrast, no enrichment for the other predicted RNA binding sites was observed, including the one in *nad2* intron 2, which did not support a direct role of MTSF3 in the splicing of this intron (Figure 6C).

**Figure 6.**
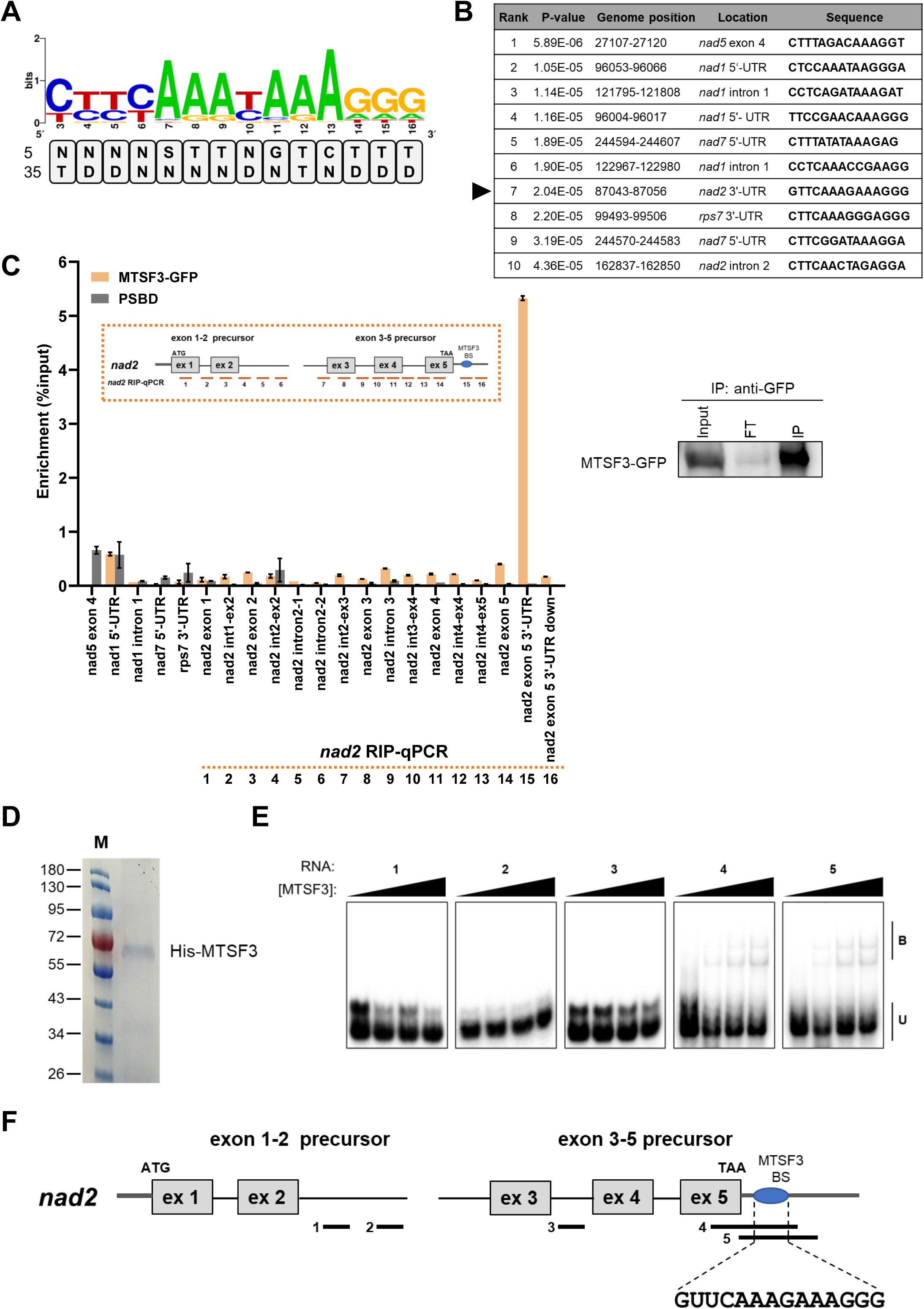
The MTSF3 protein binds specifically to the 3’ region of the *nad2* mRNA. (A) Putative binding sites of MTSF3 were predicted based on the amino acid residues at positions 5 and 35 of PPR motifs 3 to 16. Repeats are listed from N to C-terminus. The obtained combinations were then used to calculate the probabilities of nucleotide recognition by each individual PPR repeat according to the PPR code. The sequence logo depicting these probabilities was obtained with http://weblogo.berkeley.edu/. (B) Prediction of potential MTSF3 binding sites within the Arabidopsis Col-0 mitochondrial genome. The ten most-probable MTSF3 binding sequences are shown. The arrowhead marks the site located in the 3’ region of *nad2*, shown in subsequent experiments to be the RNA target of MTSF3. The *P*-values were determined with the FIMO program. (C) The MTSF3 protein specifically associates with the 3’ end extremity of the *nad2* mRNA *in vivo*. Total extracts from transgenic Arabidopsis cells expressing an *MTSF3*-GFP fusion and control untransformed PSB-D cells were used for immunoprecipitation with an anti-GFP antibody. Coimmunoprecipitated RNAs were analyzed by RT-qPCR using primer pairs of the indicated mitochondrial transcript regions. Amplification products covering *nad2* precursor transcripts are shown in the orange box above the histogram. *nad2* exon 5 3’-UTR (PCR #15) shows the enrichment obtained with a primer pair targeting *nad2* 3’ extremity. Immunoblot result of total extracts (Input), flow-through (FT) and immunoprecipitated (IP) fractions using the GFP antibody is presented. (D) SDS protein gel stained with Coomassie blue showing the purity of the HIS-MTSF3 fusion protein overexpressed and purified from *Escherichia coli*. M: protein size marker (kDa). (E) Electrophoretic mobility shift assay showing the association of MTSF3 to the 3’ region of *nad2 in vitro*. EMSA assays were performed with purified HIS-MTSF3 and the RNA probes depicted in panel F. For each probe, the reactions contained (from left to right) 0, 200, 400 and 800 nM of pure HIS-MTSF3 protein, 100 pM of gel-purified RNA probe and 2 mg*/*ml heparin. U: unbound probe; B: bound probe. (F) Diagram of the *nad2* mRNA with the relative positions of the RNA probes (numbered from 1 to 4) used in EMSA assays. The MTSF3 binding site is indicated as MTSF3 BS.

To confirm the direct association of MTSF3 with the 3’ end of *nad2* mRNA, electrophoretic mobility shift assays (EMSA) were performed. Recombinant MTSF3 lacking its mitochondrial targeting sequence was expressed in fusion with the 6XHis tag in *Escherichia coli* (Figure 6D). Five different radiolabeled RNA probes in the 3’ terminus, the intron 2 and the intron 3 regions of *nad2* mRNA were then tested for their ability to associate with the MTSF3 protein (Figure 6E). A shift was only observed for the RNA segments covering the predicted binding sequence GUUCAAAGAAAGGG located in *nad2* 3’ UTR. No binding was detected for the RNA probes contained in intron 2 or intron 3 (Figure 6E).

Additional evidence supporting an association between MSTF3 protein and the 3’ UTR of *nad2* comes from the analysis of mitochondrial small RNA species (sRNAs). Effectively, the binding of RNA-stabilizing factors such as PPR proteins was shown to lead to the accumulation of short RNA sequences in plant organelles corresponding to their binding sites (Zhelyazkova et al., 2011; Ruwe and Schmitz-Linneweber, 2012; Haili et al., 2013; Ruwe et al., 2016; Wang et al., 2017). These RNA footprints can be identified in large RNA sequence libraries produced from high-throughput small RNA sequencing programs (Ruwe et al., 2016). We thus searched the clustered organellar sRNA (cosRNA) database to see if any RNA footprints mapped to the 3′ region of *nad2* mRNA. Indeed, a 36-nucleotide RNA fragment whose 3′-extremity coincides with the MTSF3 binding site can be identified, strongly supporting an association of MTSF3 with this RNA region *in vivo* (Supplemental Figure S4). This 3’ extremity also corresponds with the predominant 3’ end of *nad2* (Forner et al., 2007).

Taken together, our *in vivo* and *in vitro* data clearly demonstrated that MTSF3 protein binds to the 3’ extremity of *nad2* mRNA in a sequence-specific manner to stabilize the 3’-end of both *nad2* mature mRNA and that of its exon 3-5 precursor.

## DISCUSSION

### MTSF3 is required for the stabilization but not the splicing of the mitochondrial *nad2* mRNA, as previously proposed

A functional characterization of the same PPR gene (called *emb2794* in this analysis) using similar approaches was recently undertaken (Marchetti et al., 2020). The author made similar observations to ours supporting that (1) the EMB2794/MTSF3 is a mitochondria-targeted PPR protein, (2) *emb2794/mtsf3* mutants are deficient in respiratory complex I because of a lack of mature *nad2* mRNA production. However, they could not firmly conclude on the function of EMB2794/MTSF3 as their RNA binding analyses uncovered many potential RNA targets, with some unrelated to *nad2*. They however favored a role of EMB2794/MTSF3 in the splicing of *nad2* intron 2 and 3, without excluding a possible function in *nad2* mRNA though (Marchetti et al., 2020).

In this analysis, we precise the function of a mitochondria-targeted PPR protein that we renamed Mitochondrial Stability Factor 3 (MTSF3) and demonstrate that it is essential for the stabilization of the mature *nad2* mRNA and, concomitantly, that of the *nad2* precursor transcript bearing exons 3 to 5, as both RNAs share the same 3’ extremity (Figure 5C). The production of the *nad2* mature mRNA effectively requires a *trans*-splicing reaction involving its second intron, allowing the five *nad2* exons to be reassembled into a single mRNA molecule. The proposed function for MTSF3 was drawn from several concordant observations. Firstly, analysis of mitochondrial transcripts showed that mature *nad2* mRNA as well as the precursor transcript carrying exon 3 to 5 strongly under-accumulate in *mtsf3* mutants. We could also observe that the *trans*-splicing reaction involving intron 2 is reduced as mature mRNAs with spliced intron 2 under-accumulate in the mutants and pre-mRNAs containing the 5’ half of intron 2 are over-abundant. However, pre-mRNAs bearing the 3’-half of *nad2* intron 2 behave inversely and slightly under-accumulate in *mtsf3* plants. Such opposed abundances of the two precursors flanking *nad2 trans-*intron 2 does not support a direct role of MTSF3 in the splicing of this intron but rather that the destabilization of the precursor carrying exon 3-5 results in the loss of the 3’ partner of *trans-*intron 2, which therefore can no longer splice in *mtsf3* mutants. The role of MTSF3 in *nad2* mRNA stabilization is also strongly supported by the RNA binding property of MTSF3 which shows high affinity to the last 14 nucleotides of mature *nad2* mRNA, according to both our *in vivo* and *in vitro* analyses (Figure 6). Moreover, a short RNA footprint matching with both the MTSF3 binding site and the 3’ end of *nad2* mRNA further argue in favor of a role of MTSF3 in *nad2* stabilization (Figure S4).

Our co-immunoprecipitation assay did not reveal any association of MTSF3 with any regions other than the 3’ end of *nad2* (Figure 6C). In particular, we did not reveal any affinity of MTSF3 for predicted RNA binding sites in the 5’ half of intron 2 or in intron 3 by gel shift assay (Figure 6D). A role of MTSF3 in *nad2* splicing is thus highly unlikely according to our results. The loss of MTSF3 induces the degradation of *nad2* exon 3-5 pre-mRNA which secondarily impacts the splicing of *nad2 trans*-intron 2 by elimination of its 3’-half partner. The impact of the *mtsf3* mutation on *nad2* intron 2 splicing is thus indirect and not direct as proposed in (Marchetti et al., 2020). Sequence analysis further revealed that MTSF3 protein in a conserved PPR protein in angiosperms like its RNA binding target at the end of *nad2* transcript (Supplemental Figure S5). Our analysis of MTSF3 reveals thus a new factor involves in plant mitochondrial mRNA stabilization and further demonstrate the importance of PPR proteins in RNA stability in plant mitochondria (Haili et al., 2013; Wang et al., 2017). These factors have in common to bind in mRNA 3’ regions and by doing so likely act as roadblock to prevent the progression of 3’-to-5’ exonucleases (*i*.*e*. the PNPase) and thereby concomitantly stabilize and define the 3’-end extremity of their target mRNA.

### *mstf3* is one of the rare examples of embryo-lethal complex I mutant

Numerous mutants affected in mitochondria-targeted PPR proteins or other kinds of RNA binding factors have been characterized, mostly in Arabidopsis, maize and rice (reviewed in (Colas des Francs-Small and Small, 2014)). In most cases, these mutants are defective in the expression of a single (or very few) mitochondrial gene(s). If we exclude the ones showing no growth alterations, most mutants fall into two major phenotypical classes corresponding either to slow growing plants or to germination defective/embryo-lethal (EL) mutants (Colas des Francs-Small and Small, 2014). The first mutant class concerns almost exclusively complex I deficient plants whose loss was found to be non-lethal to plants, whereas EL mutants tend to be affected in latter stages of the respiratory chain, like in complex III or IV, or in cytochrome *c* biogenesis (Dahan et al., 2014; Nguyen et al., 2021). Nevertheless, very few examples of mutants affected in these late stages of the plant respiratory chain were characterized up to now and the embryo-lethality associated with the *emb2794* (*mtsf3*) mutation led us to consider that it could reveal a new interesting case. Our analysis confirmed that the *mtsf3* mutants are effectively unable to germinate in normal conditions and that this deficiency is associated with an early arrest of embryo development at the late torpedo stage (Figure 1). We succeeded in repairing the embryo lethality of *mtsf3* by partial complementation with the seed specific promoter *ABI3* and could observe that rescued *mtsf3* mutants displayed a general slow-growth phenotype with curly leaves, a phenotype that is typical of Arabidopsis complex I mutants (Figure 2). Biochemical analysis confirmed that *mtsf3* mutants indeed correspond to complex I-deficient plants and that no other respiratory chain complexes are either missing or reduced in these plants. Therefore, *mtsf3* represents one of the very first examples of Arabidopsis embryo-lethal mutants deficient in complex I. What makes the *mtsf3* knock-out mutant EL as opposed to most other Arabidopsis complex I mutants? The difference of phenotype cannot be explained by the mRNA target of MTSF3, as most other Arabidopsis mutants affected in *nad2* expression were not found to be embryo-lethal (Liu et al., 2010; Kuhn et al., 2011; Hsu et al., 2014; Weissenberger et al., 2017; Wang et al., 2018). Two other Arabidopsis complex I mutants, called *msp1* and *otp43*, that are deficient in the splicing of the *nad1* mitochondrial mRNA also revealed to be EL (de Longevialle et al., 2007; Best et al., 2019). The difference of phenotypic expression between embryo-lethal complex I mutants and those that can germinate may depends on (1) the terminal developmental stage reached by mutant embryos and/or (2) the severity of complex I reduction in the different mutants (these two criteria may be directly connected). It was recently shown that the severity of vegetative growth defects in complex I Arabidopsis mutants correlates with the residual levels of complex I left in the plants (Kühn et al., 2015). Mutants with undetectable complex I levels compensate for ATP production deficit by increasing substrate level respiratory fluxes which seems somehow deleterious to plant growth. Embryo-lethal complex I mutants may represent extreme cases with harmful respiratory compensations leading to embryo developmental arrest. Alternatively, EL complex I mutants could simply accumulate insufficient levels of residual complex I, whereas non-EL mutants could contain enough complex I to produce germination competent embryos and thus escape strict embryo lethality. The connection between embryo lethality and residual complex I accumulation is currently hard to evaluate as the level of complex I in mutants has been determined by in-gel activity staining, a technique that is far from being accurate and also poorly sensitive. Using this approach, quite a few Arabidopsis mutants were shown to contain similarly low levels of complex I but they display quite different developmental alterations which should be explained by their residual complex I content (Kühn et al., 2015; Ligas et al., 2019). Our study confirms that the generally accepted non-essential character of complex I for embryo development in Arabidopsis should be re-evaluated using a more quantitative approaches to compare residual complex I accumulation in EL and non-EL complex I mutants.

## MATERIALS AND METHODS

Oligonucleotides used in this study are listed in Supporting Information Table S1.

### Plant materials

Arabidopsis (*Arabidopsis thaliana*) Col-0 plants were obtained from the Arabidopsis stock centre of the Institut National de Recherche pour l’Agriculture, l’Alimentation et l’Environnement in Versailles (http://dbsgap.versailles.inra.fr/portail/). T-DNA insertion line *mstf3* (N816708) mutants was obtained from the European Arabidopsis Stock Center (NASC) (McElver et al., 2001). The *mtsf3* mutant plants were genotyped by PCR using the primers listed in Supporting Information Table S1 and the insertion site was confirmed by sequencing. No homozygous lines could be identified in the progeny of self-fertilized heterozygous plants due to the essential role of MTSF3, as previously reported (Meinke et al., 2008; Meinke, 2020).

### Functional complementation of *mtsf3* mutants

For the partial complementation, a 2.5 kb fragment of the *ABI3* promoter (AT3G24650) was PCR amplified and inserted into the Gateway pGWB13 expression vector (Nakagawa et al., 2007) at the *Hind*III site to create p*ABI3*-GWB13 plasmid. Subsequently, the full-length coding sequence of *MTSF3* was PCR amplified and cloned into vector p*ABI3*-GWB13 by Gateway™ cloning (Invitrogen). For the full complementation assay, the *MTSF3* coding sequence but deprived of its stop codon was transferred by Gateway™ cloning (Invitrogen) into pGWB5 (Nakagawa et al., 2007) resulting in a C-terminal fusion with the GFP coding sequence, expressed under the control of the 35S promoter. The resulting constructs were then transferred into *Agrobacterium tumefaciens* strain C58C51 and introduced into heterozygous *mtsf3* plants by the floral dip method (Clough and Bent, 1998). The complemented homozygous mutant plants were identified in the progenies of generated transgenic plants.

### Subcellular localization

In this approach, the MTSF3::GFP fusion construct (see above for details) was used to transform the PSB-D *Arabidopsis* cell suspension culture as indicated in (Van Leene et al., 2015). Prior to observation, transgenic cells were soaked in 0.1 μM Mitotracker™ Red (Invitrogen) to label mitochondria. The GFP fluorescence was visualized in the transgenic cell lines by Leica TCS SP5 confocal microscopy.

### RNA extraction and analyses

Total RNAs were extracted from 8-week-old flower buds using the TRIzol reagent (Life Technologies) according to the manufacturer’s instructions. For RNA gel blotting, 10 μg of total RNA was electrophoretically separated in formaldehyde-containing (1.5% (w/v)) agarose gels and transferred onto nylon membranes (Genescreen) as described previously (Haili et al., 2013). Hybridization probes were generated by PCR amplifi cation using gene specifi c primers (Table S1) and radiolabeled with the Prime A Gene labeling kit (Promega) following the recommendations of the manufacturer.

### Quantitative RT-PCR

RNA samples were treated with DNase Max (QIAGEN) for 30 min at 37°C and transcribed using the M-MLV-Reverse Transcriptase (ThermoFisher Scientific) with random hexamer primers according to the manufacturer’s instructions. Mitochondrial transcriptome analysis and the quantification of splicing efficiency of mitochondrial introns were performed by quantitative RT-PCR as previously described (Koprivova et al., 2010; Wang et al., 2018). Two biological and three technical repeats were performed and the nuclear 18S ribosomal RNA gene was used for data normalization.

### Blue native gel and respiratory complex in-gel activity assay

Crude Arabidopsis mitochondrial extracts from eight-week-old flower buds were prepared as previously described in (Dahan et al., 2014). An equivalent to 100 mg of mitochondrial proteins were loaded and separated on 3–12% (w/v) polyacrylamide NativePAGE Bis/Tris gels (Invitrogen). Following electrophoresis, BN-PAGE gels were stained with Coomassie Blue or in buffers revealing activities of mitochondrial respiratory complexes I and IV as previously described (Sabar et al., 2005). When colorations were sufficient, gels were transferred to a fixing solution containing 30% (v/v) methanol and 10% (v/v) acetic acid to stop the reactions.

### Protein extraction and immunoblotting

Total proteins were extracted from crude membrane in cold lysis buffer (50 mM Tris-HCl, pH 7.5, 100 mM NaCl, 5 mM EDTA, 1% NP-40, 0.5% sodium deoxycholate, 0.1% SDS, 1X of complete EDTA-free protease inhibitor (Roche)). Protein concentrations were measured with Bradford (Bio-Rad), resolved on SDS-PAGE, and transferred onto PVDF membrane under semidry conditions. For BN-PAGE gel analysis, transfers were done overnight at 20 V and 4°C to PVDF membranes in transfer buffer (50 mM Bis/Tris and 50 mM Tricine). Membranes were probed with the primary antibodies listed in Supporting Information Table S2. Hybridization signals were revealed using enhanced chemiluminescence reagents (Western Lightning Plus ECL, Perkin Elmer).

### RNA Coimmunoprecipitation

RNA immunoprecipitation (RIP) assays were performed with the μMACS GFP-Tagged Protein Isolation Kit (Miltenyi Biotec) according to manufacturer’s instructions with minor modifications. Arabidopsis cells expressing the MTSF3::GFP translational fusion were collected from a 3-day-old culture and ground into a fine powder in liquid nitrogen. Samples were homogenized in RIP lysis buffer (20 mM HEPES-KOH, pH 7.6, 100 mM KCl, 20 mM MgCl_2_, 1 mM DTT, 1% Triton X-100, 1X of complete EDTA-free protease inhibitor (Roche)) for 30 min at 4°C with slow rotation (10 rpm). The lysates were clarified by centrifugation at 100,000 *g* for 20 min at 4°C and resulting the supernatant was incubated with 50 μL of anti-GFP magnetic beads (Miltenyi Biotec) for 30 min at 4°C with rotation (10 rpm). Following three washes with RIP washing buffer (lysis buffer with 0.1% Triton X-100) and recovery of the beads on a magnetic stand each time, RNAs were extracted from the beads with TRIzol reagent (Life Technologies), precipitated with ethanol, and then used for RT-qPCR analysis. Prior to retrotranscription, immunoprecipitated RNAs were treated with DNase Max (QIAGEN). Two biological and three technical repeats were performed and a RIP experiment with untransformed PSBD cells was used as negative control.

### Recombinant MTSF3 production

The *MTSF3* coding sequence deprived of the upstream region encoding the mitochondrial presequence was PCR amplified using the GWPPR-P11F2 and GWPPR-P11R2 primers (Table S1). The obtained amplification product was cloned by Gateway™ BP reaction into pDONR207 (Invitrogen) and subsequently transferred into the pDEST17 expression vector by LR reaction (Invitrogen). The 6xHis-MTSF3 fusion protein was expressed in Rosetta™ (DE3) *Escherichia coli* cells by inducing log phase cultures with 0.5 mM of isopropyl b-D-thiogalactoside at 20°C for 18 h. Following expression, the 6xHis-MTSF3 protein was solubilized from *E. coli* inclusion bodies in 150 mM NaCl, 50 mM Tris (pH 7.5) and 10% N-lauryl sarcosine, 0.5 mg/ml leupeptin and 0.06 mg/ml pepstatin. The obtained protein solution was subsequently dialyzed overnight against the same buffer in which the N-lauryl sarcosine concentration was changed to 0.01%. The protein was then enriched to high purity using a Ni-Sepharose column (GE Healthcare) and stored at 4°C before use.

### Electrophoretic Mobility Shift Assay

EMSA experiments were performed as previously described (Wang et al., 2017). Succinctly, radiolabeled RNA probes were generated by *in vitro* transcription using PCR products generated with primers listed in Supporting Information Table S1. Prior to EMSA assay, 100 pM of purified oligonucleotides were heated to 95°C for 1 min and snap-cooled on ice. Labeled RNAs were incubated with increasing MTSF3 concentrations in a buffer containing 50 mM HEPES-KOH, PH 7.8, 100 mM NaCl, 0.1 mg/ml BSA, 4 mM DTT, 10% glycerol, 2 mg/ml heparin, 10 U Riboblock at 30°C for 30 min. Reactions were then separated on 5% polyacrylamide gels containing 1 × THE (66 mM HEPES, 34 mM Tris (pH 7.7), 0.1 mM EDTA) buffer. After electrophoresis, the gels were dried and exposed to a phosphorimager screen (FLA-9500 Fujifilm).

### Bioinformatic prediction of MTSF3 binding sites

The putative RNA binding motif for MTSF3 was generated as indicated in (Wang et al., 2017). In brief, the identities of amino acid residues at 5 and 35 were extracted from each MTSF3 PPR repeat to assign a nucleotide preference according to (Barkan et al., 2012; Yan et al., 2019). The obtained nucleotide motifs were then used to search putative MTSF3 binding sites in the entire Col-0 mitochondrial genome (JF729201.1) with the FIMO program.

## ACKNOWLEDGMENTS

This work was supported by Agence Nationale de la Recherche (ANR) Grant ANR-16-CE11-0024-01. The Institut Jean-Pierre Bourgin’s (IJPB’s) benefits from the support of Saclay Plant Sciences Grant ANR-17-EUR-0007. This work has benefited from the support of IJPB’s Plant Observatory technological platforms.

## AUTHORS’ CONTRIBUTIONS

H.M. and C.W. designed research; C.W., L.B., M.Q. and C.D-G. performed research; C.W. and H.M. analyzed data; C.W. and H.M. wrote the paper.

## SUPPORTING INFORMATION

Additional Supporting Information may be found in the online version of this article.

**Supplemental Figure S1. The *MTSF3* gene encodes a mitochondria-targeted PPR Protein**. Confocal microscope images showing the subcellular distribution of a MTSF3::GFP translational fusion in transgenic Arabidopsis PSB-D cells. (Left) GFP fluorescence. (Center) MitoTracker™ fluorescence. (Right) Merged signals. Scale bars, 26.9 μm.

**Supplemental Figure S2. BN-PAGE analysis of the respiratory chain complexes in p*ABI3*::*MTSF3* plants**. (A) Crude mitochondrial extracts prepared from wild-type and two p*ABI3*::*MTSF3* plants (#1 and #2), separated on BN-PAGE gels and stained with Coomassie blue. In-gel staining revealing cytochrome *c* oxidase activity of complex IV are also presented. (B) Following migration, BN-PAGE gels were transferred to membranes and blots were hybridized with antibodies to mitochondrial RISP and ATPβ. The different respiratory complexes are designated by their roman numeral.

**Supplemental Figure S3. The alternative respiratory pathway is induced in p*ABI3*::*MTSF3* mutant plants**. Quantitative RT-PCR results measuring the relative accumulation levels of alternative oxidase (*AOX*) and NADH dehydrogenase (*NDA, NDB* and *NDC*) transcripts in p*ABI3*::*MTSF3* mutant and wild type plants. Two biological repeats and three technical repeats were performed for each genotype in this analysis.

**Supplemental Figure S4. A short RNA footprint corresponding to the 3’ end of *nad2* transcripts and overlapping with the MTSF3 binding site accumulate *in planta***. The diagrams show read coverage of a clustered organellar short RNA (cosRNA) mapped to the 3’ regions of the *nad2* transcript. This cosRNA (M86) is shown as blue box and the minimal binding site of MTSF3 is marked as dotted line. The shown data were extracted from https://www.molgen.hu-berlin.de/projects-jbrowseathaliana.php, which reports identified cosRNAs in Arabidopsis plastid and mitochondrial genomes. The two red arrows correspond to the predominant 3’ ends of *nad2* mRNA, according to (Forner et al., 2007).

**Supplemental Figure S5. The sequence and the position of MTSF3 binding site are conserved in angiosperms**. (A) Multiple sequence alignment of Arabidopsis MTSF3 protein with putative orthologous proteins from a representative selection of dicot (*Brassica rapa, Erythranthe guttata*) and monocot (*Oryza sativa, Zea mays*) species. The extent of each PPR repeats is indicated with red lines. Residue 5 and 35, which have been shown to be essential for PPR protein-RNA specific interaction, are marked with red and green asterisks, respectively. The grey bar represents MTSF3 mitochondrial targeting region, as predicted by TargetP. (B) Predicted RNA binding sites of Arabidopsis MTSF3 and its orthologous proteins from the indicated plant species. Five and 35 amino acid combinations are listed from N to C-terminus. The obtained combinations were then used to calculate the probabilities of nucleotide recognition by each individual PPR repeat according to the PPR code. The sequence logo depicting these probabilities was obtained with http://weblogo.berkeley.edu/. (C) Multiple sequence alignment of the 3’ UTR of *nad2* from the above indicated plant species. The cosRNA (M86) corresponding to the MTSF3 binding site is shown as green bar and the minimal binding site of MTSF3 is marked as dot line.

**Table S1**. List of primers used in this study.

**Table S2**. List of antibodies used in this study.

